# Enteric Glial Cell Network Function is Required for Epithelial Barrier Restitution following Intestinal Ischemic Injury in the Early Postnatal Period

**DOI:** 10.1101/2022.11.04.514575

**Authors:** Amanda L. Ziegler, Sara Erwin, Madison L. Caldwell, Melissa S. Touvron, Tiffany A. Pridgen, Scott T. Magness, Jack Odle, Laurianne Van Landeghem, Anthony T. Blikslager

## Abstract

Ischemic damage to the intestinal epithelial barrier, such as in necrotizing enterocolitis or small intestinal volvulus, is associated with higher mortality rates in younger patients. We have recently reported a powerful pig model to investigate these age-dependent outcomes in which mucosal barrier restitution is strikingly absent in neonates but can be rescued by direct application of homogenized mucosa from older, juvenile pigs by a yet-undefined mechanism. Within the mucosa, a postnatally developing network of enteric glial cells (EGC) is gaining recognition as a key regulator of the mucosal barrier. Therefore, we hypothesized that the developing EGC network may play an important role in coordinating intestinal barrier repair in neonates. Neonatal and juvenile jejunal mucosa recovering from surgically induced intestinal ischemia was visualized by scanning electron microscopy and the transcriptomic phenotypes were assessed by bulk RNA sequencing. EGC network density and gliosis were examined by gene set enrichment analysis, three-dimensional volume imaging and western blot and its function in regulating epithelial restitution assessed *ex vivo* in Ussing chamber using the glia-specific inhibitor fluoroacetate, and *in vivo* by co-culture assay. Here we refine and elaborate our translational model, confirming a neonatal phenotype characterized by a complete lack of coordinated reparative signaling in the mucosal microenvironment. Further, we report important evidence that the subepithelial EGC network changes significantly over the early postnatal period and demonstrate that EGC function in close proximity to wounded intestinal epithelium is critical to intestinal barrier restitution following ischemic injury.

**NEW & NOTEWORTHY:** This study refines a powerful translational pig model, defining an age-dependent relationship between enteric glia and the intestinal epithelium during intestinal ischemic injury and confirming an important role of the enteric glial cell activity in driving mucosal barrier restitution. This study suggests that targeting the enteric glial network could lead to novel interventions to improve recovery from intestinal injury in neonatal patients.

## INTRODUCTION

The intestinal epithelium is sensitive to a range of digestive disease insults, particularly those characterized by ischemia, because of the villus counter-current blood flow that exacerbates epithelial hypoxia at the tips of villi.(1, 2) Ischemia-induced breakdown of the intestinal epithelial barrier leads to sepsis, intestinal necrosis and host death unless the barrier is rapidly and completely repaired following an ischemic event.(1-4) Intestinal ischemia is a component of devastating diseases such as strangulating obstruction and neonatal necrotizing enterocolitis.(5) For example, a cause of strangulating obstruction in infants, volvulus, most commonly occurs because of anatomical mesenteric abnormalities and, in terms of age, has the highest reported mortality in the immediate newborn period.(6) Despite many advances in the management of critical patients with gastrointestinal failure, severe intestinal injuries which impact the intestinal barrier persistently have much worse reported clinical outcomes in younger patient populations. A disease associated with particularly severe barrier dysfunction, necrotizing enterocolitis, is associated with unremittingly high mortality rates between 20 and 30%, with the youngest patients, preterm neonates, suffering the worst outcomes.(7, 8) These poorer clinical outcomes in neonates as compared to older patients have not been fully explained, and this represents a critical limitation to our ability to improve survival in these patients.

Clinical interventions for neonatal intensive care unit patients with ischemia-injured intestine currently include surgical correction of the obstruction, supportive care, attention to sepsis that results from disruption of the intestinal barrier, and ultimately intestinal resection when necessary.(9, 10) Novel treatments have focused on stimulating *de novo* formation of new epithelial cells from the stem cell niche, but this requires support of the patient for days following the initial injury until newly produced epithelial cells can restore the mucosal barrier.(9, 11) In the early reparative phase (within hours), remaining epithelium can immediately restitute the damaged barrier to curtail sepsis early on in the clinical onset of disease and prevent patient mortality until the regenerative phase can fully restore intestinal architecture.(1) For this reason, our lab has focused on understanding the mechanisms of the early phase of repair, as clinical interventions enhancing this acute reparative phase are particularly promising for improving patient survival.

We study mechanisms of acute mucosal repair following ischemia using a porcine model because pigs are widely appreciated as one of the most powerful comparative models for human biology and disease, and for gastrointestinal disease in particular, due to many important similarities between porcine and human diet, size, and gastrointestinal physiology.(12-17) The vascular anatomy of humans and pigs has been shown to be remarkably similar, with comparable levels of epithelial injury induced by oxidative enzymes and neutrophil infiltration during ischemic events.(2, 18) In juvenile (6-8-week-old) pigs, we have shown rapid restitution and recovery of intestinal barrier function, effectively minimizing sepsis and complications in the host after injury. We recently found that in neonatal (2-week-old) pigs the epithelial barrier failed to restitute under the same conditions.(19) An age-dependent defect in epithelial restitution in neonates is likely to be a critical contributor to the high morbidity and mortality seen in infant patients clinically affected by intestinal ischemia and barrier injury. Therefore, this newly identified age-dependent intestinal repair defect in a highly comparative porcine model can provide critical insight for addressing the unacceptably high morbidity and mortality rates suffered by the youngest of critically ill patients. Importantly, we have found that neonatal restitution can be rescued by direct application of ischemic-injured homogenized mucosa from juvenile pigs, suggesting that neonatal epithelial cells are intrinsically capable of efficient restitution in response to yet uncharacterized cues present in juvenile mucosa.(19) Therefore the identification of the components of the juvenile mucosal tissues which rescue neonatal restitution will inform future development of novel interventions for devastating intestinal diseases associated with age-dependent defects in barrier repair mechanisms.

A complex and dense population of mesenchymal, circulatory, immune, and neural cells within the lamina propria are known to modulate epithelial cell functions in homeostasis and disease states by secreting paracrine factors to orchestrate complex, coordinated signaling in the mucosal microenvironment. Work in rodents suggests that some of these cell populations are fully developed at birth, whereas others, particularly enteric glia, develop postnatally to form a dense network in close proximity to intestinal epithelial cells. This glial network continues maturation prior to and following weaning during the postnatal period in rodents.(20-22) Additionally, a growing body of evidence indicates that enteric glial cells play a pivotal role in promoting intestinal epithelial barrier function and healing through paracrine signaling mechanisms(23-28). This led us to hypothesize that the developing glial network may play an important role in the establishment of efficient mechanisms of intestinal barrier repair during the postnatal period. In this study, we sought to examine how the subepithelial enteric glial network changes over the early postnatal period, and how this impacts intestinal barrier restitution following ischemic injury using our porcine model.

## MATERIALS & METHODS

### Animals

All procedures were approved by NC State University Institutional Animal Care and Use Committee. Commercial crossbred pigs 2- or 6-weeks-of-age of either sex were euthanized at an academic animal production facility for immediate tissue collection or were transported to a large animal surgical facility for surgical modeling experiments.

### Surgical Ischemic Injury Model

Pigs were sedated using xylazine (1.5 mg/kg) and ketamine (11 mg/kg). Anesthesia was induced with isoflurane vaporized in 100% oxygen via face mask, after which pigs were orotracheally intubated for continued delivery of isoflurane to maintain general anesthesia. Pigs were placed on a water-circulated heating pad and intravenous fluids were administered at a maintenance rate of 15 ml·kg^-1^·h^-1^ throughout surgery. The distal jejunum was accessed via midline or paralumbar incision and 8-10 cm loops were ligated in segments and subjected to 30-minutes of ischemia via ligation of local mesenteric blood vessels with 2-0 braided silk suture. Adjacent loops not subjected to ischemia were used as control tissue. For enteric glial cell inhibition studies, Ringers solution containing 0μM, 5μM, 500μM or 5000μM sodium fluoroacetate (MP Biomedicals Cat #ICN201080) was injected into the lumen of the intestinal loops. At the time of tissue collection, pigs were euthanized with an overdose of pentobarbital. Intestinal loops were excised and opened longitudinally along the antimesenteric border and placed in cold, oxygenated Ringers solution. For enteric glial cell inhibition studies, tissues were transported from the surgical facilities to the laboratory for Ussing chamber experiments in cold, oxygenated Ringers solution containing 0μM, 5μM, 500μM or 5000μM sodium fluoroacetate.

### Scanning electron microscopy

30-minute ischemic neonate and juvenile jejunum were fixed at 30- and 120-minutes during the ex vivo recovery period in independent experiments. Mucosa was rinsed briefly with PBS to remove surface debris followed by immersion fixation in 2% paraformaldehyde/2.5% glutaraldehyde/0.15M sodium phosphate buffer, pH 7.4. Specimens were stored in the fixative overnight to several days at 4°C before processing for SEM (Microscopy Services Laboratory, Dept. of Pathology and Laboratory Medicine, UNC, Chapel Hill, NC, USA). After three washes with 0.15M sodium phosphate buffer (PBS), pH 7.4, the samples were post-fixed in 1% osmium tetroxide in PBS for 1-hour and washed in deionized water. The samples were dehydrated in ethanol (30%, 50%, 75%, 100%, 100%), transferred to a Samdri-795 critical point dryer and dried using carbon dioxide as the transitional solvent (Tousimis Research Corporation, Rockville, MD). Tissues were mounted on aluminum planchets using silver paste and coated with 15nm of gold-palladium alloy (60Au:40Pd, Hummer X Sputter Coater, Anatech USA, Union City, CA). Images were taken using a Zeiss Supra 25 FESEM operating at 5kV, using the SE2 detector, 30μm aperture, and a working distance of 10 to 12mm (Carl Zeiss Microscopy, LLC, Peabody, MA).

### RNA Sequencing Analysis

Jejunal mucosal scrapings were snap frozen in liquid nitrogen and stored at -80°C until further processing. mRNA were isolated using the RNeasy Mini Kit (Qiagen, Hilden, Germany) and cDNA libraries were generated using SMARTer Stranded Total RNA Sample Prep Kit - HI Mammalian (Takara Bio, Inc., Kusatsu, Japan). Libraries were high-output single-end sequenced on a NextSeq500 instrument (Illumina, Inc., San Diego, CA, USA) for 75 cycles using an 8 base pair index. High quality Sequences were aligned to an annotated porcine genome (Sscrofa11.1, University of California, Santa Cruz, CA, USA). Prior to analysis all non-transcribed genes were removed and all genes that had more than one count per million in two or more samples were analyzed.

### Gene expression analysis

Using raw counts data from NextSeq500, differential gene expression analyses between neonatal and juvenile injured and uninjured mucosal scrapings were performed using R software (R Foundation for Statistical Computing, Vienna, Austria) with the DESeq2 package (Bioconductor) and RStudio software (RStudio, Boston, MA, USA) to obtain log fold change (logFC), false discovery rates (FDR) and p-values as well as generate principal component analysis plots.(29-31)

### Pathway and Gene Set Analysis

Differentially expressed genes identified by DESeq2 were analyzed with Ingenuity Pathways Analysis (Qiagen, Hilden, Germany) to define global regulation of key “Diseases and Functions” pathways related to epithelial cell migration, adhesion, and intestinal injury and barrier function. These functions included “Cell movement,” “Migration of cells,” “Cell survival,” “Activation of cells,” “Organization of cytoskeleton,” “Cellular homeostasis,” “Microtubule dynamics,” “Organization of cytoplasm,” “Inflammation of organ,” “Cell-cell contact,” “Formation of cell protrusions,” “Formation of filaments,” and “Formation of cytoskeleton.”

As a complementary approach, Gene Set Enrichment Analysis (GSEA) was used to predict pathways and functions differentially activated between the experimental groups by determining whether predefined gene sets showed significant enrichment scores when comparing expression profiles of experimental groups. “Hallmark” and “Reactome” gene sets and the following selected Gene Ontology Biological Processes (GO BP, http://geneontology.org/) gene sets were assessed: “Glial Cell Development” “Glial Cell Proliferation” “Glial Cell Differentiation” and “Glial Cell Migration” between age groups and injury groups. Finally, a gene set defining “Gliosis” reported by Schneider *et al* in 2021 was interrogated using GSEA.(32)

### Western Blot Analysis

Distal jejunum was rinsed in cold PBS and opened along the antimesenteric border. Tissue was pinned to a transparent silicone-coated glass petri dish mucosal side up under a binocular macroscope (National Optical & Scientific Instrument Inc, Schertz, Texas, USA). The mucosa to the level of the muscularis mucosa was micro-dissected, collected, and snap-frozen in liquid nitrogen until further processing. To isolate proteins, tissues were homogenized using a handheld homogenizer (Omni International, Kennesaw, GA, USA) in T-PER tissue extraction reagent (Thermo Fisher Scientific, Waltham, MA, USA, Cat #FNN0071) with Halt Protease Inhibitor Cocktail (Thermo Fisher Scientific, Cat # 87785). Samples were centrifuged at 10,000 RPM for 5 minutes at 4°C to pellet tissue debris. Supernatant was transferred to fresh tubes and the protein concentrations of the samples were measured with a Pierce BCA protein assay kit (Thermo Fisher Scientific Cat# 23227). Samples were prepared for electrophoresis by denaturing 20μg of protein per sample in XT Sample Buffer (Bio-Rad, Hercules, CA, USA, Cat# 161-0791) and XT Reducing agent (Bio-Rad Cat# 3161-0792) at 95°C for 5 minutes. Samples were loaded into XT Criterion 4-12% Bis-Tris gels (Bio-Rad Cat #345-0123) with Precision Plus Protein Dual Color Standards (Bio-rad Cat# 1610374) and run at 200V for 35 minutes in XT MES Running Buffer (Bio-Rad Cat# 161-0789). Gels were transferred to a PVDF membrane (Bio-Rad Cat# 162-0175) at 100V, 1.5A for 30 minutes in 20% methanol Tris/Glycine Buffer (Bio-Rad Cat# 161-0734). Membranes were blocked in 5% Blotting-Grade Blocker (Bio-Rad Cat# 170-6404) in tris-buffered saline buffer (Bio-rad Cat# 170-6435) with 0.01% Tween 20 (TBS/T) for 1 hour at room temperature. Membranes were incubated in the following primary antibodies overnight at 4°C: polyclonal rabbit anti-beta actin IgG (Abcam Cat# ab8227, RRID:AB_2305186) at a dilution of 1:1,000, polyclonal rabbit anti-GFAP IgG (Agilent Cat# Z0334, RRID:AB_10013382) at a dilution of 1:1,000 in 5% Blotting-Grade Blocker in TBS/T or 5% bovine serum albumin in TBS/T. Membranes were then incubated in horseradish peroxidase (HRP)-conjugated goat anti-rabbit IgG (Thermo Fisher Scientific, Cat #32460) in 5% Blotting-Grade Blocker. Blots were activated with Pierce ECL Substrate (Bio-Rad, Cat #170-5060) and imaged with a Chemidoc imaging system (Bio-rad, Cat #170-8280). Membranes were stripped between each probe with Restore PLUS Western Blot Stripping Buffer (Thermo Fisher Scientific, Cat #46430). Band densities were measured with Image J software (NIH, Bethesda, MD, USA) and normalized to β actin density as a sample loading control for quantification.

### Ussing Chamber Studies

The outer seromuscular layer was stripped by blunt dissection in cool, oxygenated Ringers solution. Stripped jejunal tissue was mounted in 1.12 cm^2^ aperture Ussing chambers. The tissues were bathed in 10 ml warmed, oxygenated (95% O_2_/ 5% CO_2_) Ringers solution on the basolateral and apical sides. For enteric glial cell inhibition studies, Ringers solution was treated with 0μM, 5μM, 500μM or 5000μM sodium fluoroacetate on both the basolateral and apical sides. Basolateral Ringers solution also contained 10mM glucose while the apical Ringers solution was osmotically balanced with 10mM mannitol. Bathing solutions were circulated in water-jacketed reservoirs and maintained at 37°C. The spontaneous potential difference (PD) was measured using Ringers-agar bridges connected to calomel electrodes, and the PD was short-circuited through Ag-AgCl electrodes with a voltage clamp that corrected for fluid resistance. Resistance (Ω·cm^2^) was calculated from spontaneous PD and short-circuit current (I_sc_). If the spontaneous PD was between -1 and 1 mV, the tissues were current-clamped at ± 100 μA and the PD re-recorded. I_sc_ and PD were recorded every 15-minutes for 120-minutes. From these measurements, transepithelial electrical resistance (TEER) was calculated.

### Light microscopy and histomorphometry

Tissues were fixed for 18 hours in 10% formalin at room temperature immediately following ischemic injury or after 120-minutes *ex vivo* recovery period. Formalin-fixed tissues were transferred to 70% ethanol and then paraffin-embedded, sectioned (5μm) and stained with hematoxylin and eosin for morphological and morphometric analyses. For morphometric analysis of villus injury, the base of the villus was defined as the opening of the neck of the crypts and height of epithelialization, total height and width of villus were measured using Image J Software (NIH, Bethesda, MD, USA). The surface area of the villus was calculated as previously described using the formula for the surface area of a cylinder modified by subtracting the base of the cylinder and adding a factor that accounted for the hemispherical shape of the villus tip (33, 34). The percentage of villus epithelialization was used as an index of epithelial injury and restitution.

### Immunofluorescence

Mucosal samples were trimmed and fixed for 18 hours in 4% paraformaldehyde (PFA) in PBS at 4°C. Fixed tissues were transferred to 10% sucrose, then 30% sucrose in PBS for 24 hours each for cryopreservation. The tissues were then embedded in optimal cutting temperature (OCT) compound and sectioned (7μm) onto positively charged glass slides for immunostaining. Sections were rehydrated 3 times in PBS before permeabilization in 0.3% Triton-X in PBS for 20 minutes. Following permeabilization, slides were washed twice in PBS then blocked (Agilent Cat# S0909) for 1 hour at room temperature. Slides were incubated overnight at 4°C in rabbit anti-GFAP (Agilent Cat# Z0334, RRID:AB_10013382) at a dilution of 1:2000 in antibody diluent (Agilent Cat# S0809). After washing slides 3 times in PBS, slides were incubated for 1 hour at room temperature in the dark in goat anti-rabbit conjugated to Alexa Fluor 647 (Molecular Probes Cat# A-21245, RRID:AB_141775) diluted 1:500 in diluent (Agilent Cat# S0809). Tissues were counterstained with the nuclear marker 4′,6-diamidino-2-phenylindole (DAPI, Invitrogen Cat# D1306) at a dilution of 1:1000 in diluent (Agilent Cat# S0809) at room temperature. Slides were washed three times in PBS then mounted under a coverslip in aqueous mounting medium (Agilent Cat# S3025). Images were captured using an inverted fluorescence microscope (Olympus IX81, Tokyo, Japan) with a digital camera (ORCA-flash 4.0, Hamamatsu, Japan) using a 20x objective lens (LUC Plan FLN, Olympus, Tokyo, Japan). One tissue per treatment was incubated with secondary antibody mixture and no primary antibody mixture serving as negative controls to ensure antibody specificity. Immunofluorescence intensity was quantified using ImageJ software. Enteric glial cells of the submucosal ganglia were identified in each individual ganglion by expression of glial fibrillary acidic protein (GFAP) co-staining with DAPI. GFAP fluorescence signal was quantified for 5 ganglia per individual (3 individuals per treatment, n =15 ganglia per treatment). Ganglia were outlined using the magic wand tool at a threshold of 18. Three background regions with no visible fluorescence were randomly selected and measured for each ganglion. Corrected total cell fluorescence (CTCF) was calculated for each ganglion as CTCF = Integrated density of glia – (Area of glia x mean gray value of background regions).

### iDISCO Volume Imaging and Quantification

EGC network was labeled and visualized in 3 dimensions using the iDISCO+ method adapted to intestinal specimens as previously published by our group.(35) Antibodies for immunolabeled three-dimensional imaging of solvent-cleared organs (iDISCO) analysis were validated for use with methanol by cryosectioning 20μm sections of PFA fixed and frozen jejunum onto glass slides, incubating the sections for 3 hours in 100% methanol, rehydrating the sections in PBS and proceeding with standard immunostaining protocol described above as recommended by the iDISCO method (https://idisco.info/idisco-protocol/). Full-thickness sections of jejunum were rinsed well in cold PBS and fixed in 4% PFA in PBS at 4°C overnight. Once fixed, tissues were trimmed to 3mm x 5mm pieces and processed for solvent clearing according to established protocols.(36) Samples were dehydrated gradually in methanol, cleared in 66% dichloromethane/33% methanol overnight at room temperature, then bleached with 5% hydrogen peroxide in methanol overnight at 4°C. Samples were then rehydrated gradually and incubated in PBS/Triton-X/glycine/dimethyl sulfoxide (DMSO) solution for 2 days at 37°C, then blocked in PBS/Triton-X/donkey serum/DMSO for 2 days at 37°C. Samples were incubated in methanol-validated polyclonal rabbit anti-GFAP IgG (Agilent Cat# Z0334, RRID:AB_10013382) at a dilution of 1:1000 in PBS/Triton-X/donkey serum/DMSO solution for 4 days at 37°C. After washing, samples were incubated in methanol-validated anti-rabbit AlexaFluor 647-conjugated secondary antibody (Thermo Fisher Scientific Cat# A-21245, RRID:AB_2535813) for 4 days at 37°C. Samples were dehydrated as above with additional 100% dichloromethane washes, and cleared in dibenzyl ether. Samples were stored in dibenzyl ether until imaging with a Lavision Ultramicroscope II light-sheet system (Lavision BioTech, Bielefeld, Germany).

Data were visualized and analyzed using Imaris Image Analysis Software (Bitplane AG, Zurich, Switzerland). To quantify subepithelial EGC density within the mucosa, five villi were randomly selected for each sample after viewing in slice mode in Imaris to ensure that the villi could be clearly visualized to the depth of the muscularis mucosa using autofluorescence of tissue landmarks. The entire villus and the area just beneath its footprint were masked manually using the click to draw surfaces feature from villus tip to the level of the muscularis mucosa. Using the surfaces feature, a surface was interpolated from these hand-drawn bounds and total villus volume was recorded from the statistics tab for each villus. Following manual villus structure isolation, the surfaces wizard was used to manually optimize an algorithm that best identified and marked the true GFAP signal in each villus. The surfaces wizard was then applied to each masked villus structure without additional adjustment to obtain signal volume measurements from the statistics tab. These values were then used to calculate percent GFAP signal by volume within the lamina propria of each villus, corresponding to the density of EGC within the lamina propria of each villus.

### EGC isolation and culture

24-well tissue culture plates were coated overnight at room temperature with 0.5mg/mL Poly-L-lysine (Sigma Cat# P2636) in 0.5M borate buffer then washed twice with sterile PBS. Jejunal layers containing either submucosal or longitudinal muscle myenteric plexus were microdissected from non-injured pigs. Tissues were next enzymatically dissociated in 1 mg/ml protease (Sigma, Cat# P4630) and 0.25mg/ml collagenase (Sigma, Cat# 9891). Following enzyme inactivation and additional mechanical dissociation, cell preparations were strained sequentially through 100μm, 70μm and 40μm cell strainers to enrich for single cells. Cells were next seeded onto fresh poly-L-lysine coated tissue culture plates and maintained in DMEM/F12 medium supplemented with 0.5% heat inactivated FBS, 1x GlutaMAX (Gibco, Cat# 35050061) 1x B27(Gibco Cat# 17504044), 1x N2 (Gibco Cat# 17502048) and 1x G5 (Gibco Cat# 17503012) supplements to enrich cultures for glial cells. Cells were passaged with 0.01% trypsin in PBS at 80% confluence. EGC were utilized for all co-culture experiments at passage 1.

### Immunocytochemistry

To assess the purity of EGC primary cultures, cells were fixed at passage 1 at approximately 50% confluence, permeabilized and saturated for 30 minutes at room temperature in PBS with 0.01% Triton-X PBS and 4% donkey serum and incubated with chicken anti-GFAP (Abcam, ab4674) and rabbit anti-S100β (Dako, z0311) antibodies diluted 1:1000 and 1: 2000 respectively, in PBS with 0.01% Triton-X PBS and 4% donkey serum. Cells were then incubated in Alexa Fluor 488 and Alexa Fluor 594-conjugated secondary antibodies against chicken (Molecular Probes Cat# A-11039, RRID:AB_142924) and rabbit (Thermo Fisher Scientific Cat# A-21207, RRID:AB_141637) at a 1:500 dilution for 2 hours at room temperature. Cell nuclei were stained with DAPI (Invitrogen Cat# D1306) at a dilution of 1:1000 in PBS for 5 mins at room temperature. Images were captured using an inverted fluorescence microscope (Olympus IX81, Tokyo, Japan) with a digital camera (ORCA-flash 4.0, Hamamatsu, Japan) using a 20x objective lens (LUC Plan FLN, Olympus, Tokyo, Japan). Color channels were merged and re-colorized in ImageJ software (NIH, USA) to visualize overlay images.

### IPEC-J2 cell scratch wound assay

IPEC-J2 cells (an intestinal epithelial cell line originally isolated from the jejunum of a neonatal pig) were cultured seeded onto Transwell inserts (Corning Cat# 3470) and maintained in DMEM/F12 supplemented with 10% heat-inactivated FBS, 5ng/mL insulin/ transferrin/ selenium supplement (Gibco Cat# 41400045) and 5 ng/mL recombinant human EGF (Gibco Cat# PHG0311). Once IPEC-J2 reached confluence, Transwell inserts were transferred to wells containing EGC primary cultures (passage 1) or blank wells containing only EGC culture medium. After 24-hours, scratch-wounds were inflicted manually with a disposable 200μL micropipette tip. Wounds were imaged at 0-, 2-, 4- and 6- hours after scratch-wounding using an inverted microscope (Olympus IX81, Tokyo, Japan) with a digital camera (ORCA-flash 4.0, Hamamatsu, Japan) using a 20x objective lens (LUC Plan FLN, Olympus, Tokyo, Japan) and plate mapping software (CellSens, Olympus, Tokyo, Japan). Percent wound healing was calculated after measuring total wound area using ImageJ software (NIH, USA).

### Statistical Analysis

Data were analyzed using Prism statistical software (Graphpad, La Jolla, California, USA). TEER and histomorphometry data were reported as means ± SEM for a given number (*n*) of animals for each experiment. Results were analyzed by unpaired Mann-Whitney test for nonparametric data, two-way ANOVA (or mixed-model on datasets with missing data points) or two-way ANOVA on repeated measures. For analyses where significance was detected by ANOVA, Sidak’s test was utilized for *post hoc* pairwise multiple comparisons. The α-level for statistical significance was set at P<0.05. CTCF data are presented as boxplots indicating the minimum, lower quartile, median, upper quartile, and maximum corrected total cell fluorescence values for each treatment group. A Wilcoxon test for nonparametric data was conducted to compare the distribution of CTCF values between the four conditions. P < 0.05 was considered significant. The Grubb’s test was used to detect and remove statistical outliers (greater than two-times standard deviations from the mean) from datasets.

### Access to Data

All authors had access to all data and have reviewed and approved the final manuscript. Complete raw Illumina FASTQ files used for RNA sequencing analyses are deposited into Gene Expression Omnibus (series entry GSE212533) for public access.

## RESULTS

### A restitution phenotype in epithelial cells fails to initiate to repair ischemia-induced mucosal wounds in neonatal intestinal mucosa

To evaluate and define the age-dependent repair phenotype of small intestinal epithelium, we examined the surface ultrastructure of epithelial wound margins in recovering neonatal and juvenile jejunum by scanning electron microscopy (Fig. 1). At 30-minutes *ex vivo* recovery (Fig. 1A, upper panels), juvenile pigs demonstrate a distinct migratory phenotype in wound-adjacent epithelial cells at the wound margins (broken line, wound bed marked with an asterisk). The epithelial cells at the wound margins exhibit flattened morphology and recruit redundant cell membrane from microvilli for incorporation into the extending lamellipodia (closed arrowheads) and filopodia (open arrowheads) as the cells migrate across the exposed basement membrane. Later in recovery (120-minutes, lower panels), the juvenile wounds are completely closed, and newly restituted epithelium, with its characteristic flattened and overlapping morphology, is visible at the villus tip (asterisk). In neonatal pigs after 30-minutes *ex vivo* recovery, a large wound bed (Fig. 1B, broken line) is surrounded by inactive wound-adjacent epithelial cells (closed arrowhead) which fail to form flattened morphology or projections characteristic of the migratory phenotype. After 120 minutes of *ex vivo* recovery (lower panels), the neonatal wound bed persists (broken line, right panel). These images reveal that morphological changes associated with restitution are absent in neonatal mucosa beyond the period of time required for juvenile mucosa to restitute effectively.

**Figure 1.**
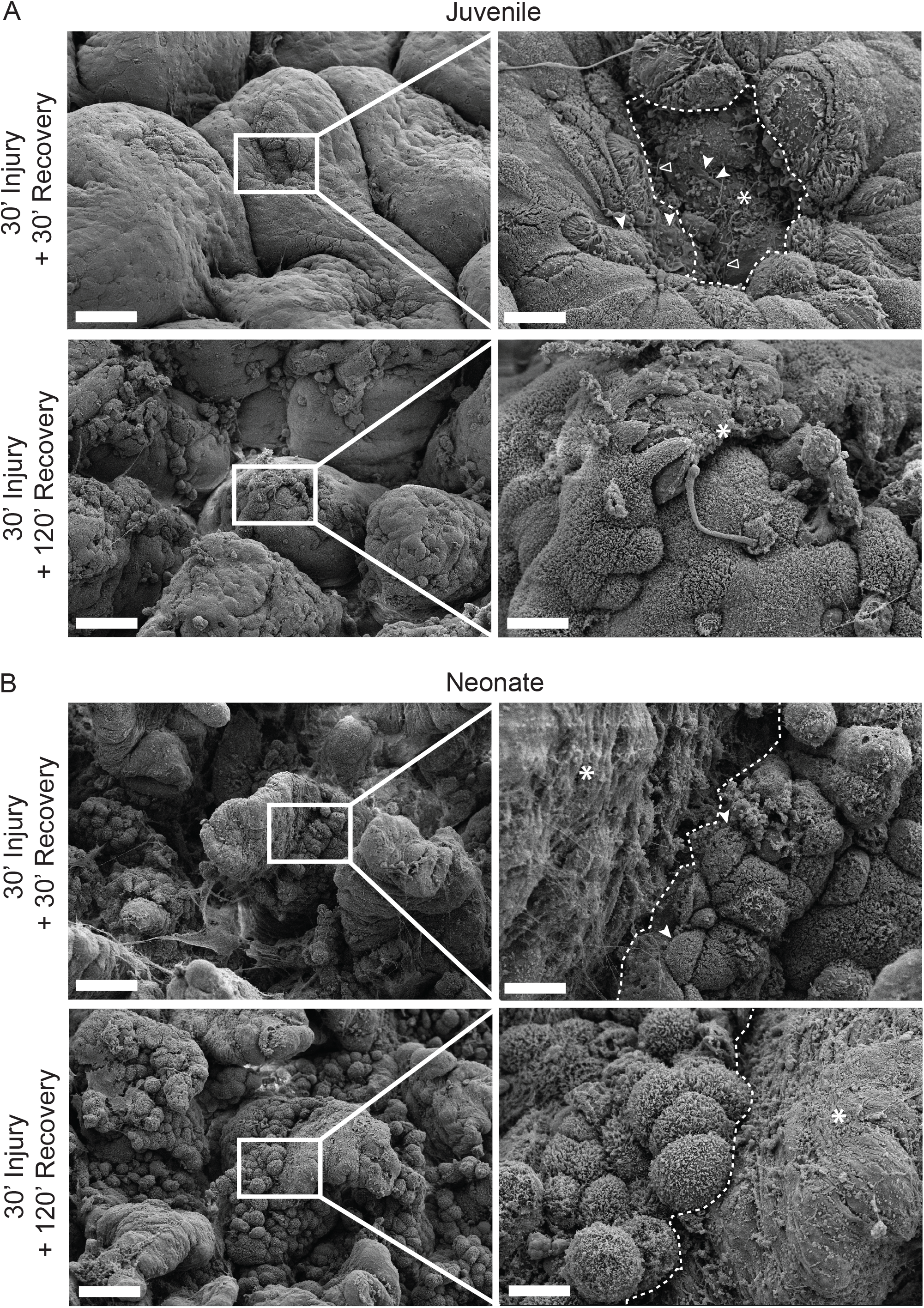
Scanning electron microscopy reveals persistent epithelial defect in neonatal pigs characterized by a lack of migratory phenotype in the wound adjacent epithelial cells observed in juveniles. (A) At 30 minutes *ex vivo* recovery (upper panels), juvenile pigs demonstrate migratory phenotype in wound-adjacent epithelial cells at the wound (asterisk) margins (broken line). Note the flattening cellular morphology and recruitment of redundant cell membrane from microvilli into the extending lamellipodia (closed arrowhead) and filopodia (open arrowhead). After 120 minutes of *ex vivo* recovery (lower panels), the wound is completely closed, and newly restituted epithelium is visible at the villus tip (asterisk). (B) At 30 minutes *ex vivo* recovery (upper panels), neonatal pigs demonstrate a large wound bed (asterisk, border marked with broken line) surrounded by inactive wound-adjacent epithelial cells (closed arrowhead) which fail to form flattened morphology or migratory projections. After 120 minutes of *ex vivo* recovery (lower panels), the wound bed persists (asterisk, border marked with broken line) and has enlarged, indicative of further lost cell adhesion and sloughing. Low magnification (left) 1000x, scale bar 40μm. High magnification (right) 5000x, scale bar 8μm.

To define this phenotype further, we assessed mucosal transcriptional profiles of the neonatal and juvenile jejunum under both ischemic and non-ischemic conditions by bulk RNA sequencing. Plotting all four groups revealed distinct age-dependent clustering and a more apparent effect of ischemia in separating the control from ischemic juveniles than control from ischemic neonates (Fig. 2 A). Plotting only the juvenile groups shows clustering of ischemic mucosal samples from control, mostly along the x-axis which accounted for 52% of the total sample variance, indicative of a coordinated transcriptional response to ischemia in juveniles (Fig. 2 B, right panel). Within neonatal mucosa, no appreciable clustering was observed when comparing control to ischemic conditions, indicative of a lack of consistent transcriptional response to ischemic injury across the three biological replicates sequenced in this age group (Fig 2 B, left panel). Plotting control tissue only showed distinct clustering of neonatal mucosal samples from juveniles along the PC1, with 68% variance along that axis (Fig. 2 C, left panel). PCA plots of just the ischemic groups demonstrate very tight clustering of ischemic juvenile mucosa distinct from ischemic neonatal mucosa on principal component analysis plots with most of the separation occurring in the x-axis accounting for 80% of the total sample variance, indicative of a differing transcriptional response in neonates as compared to juveniles under ischemic conditions (Fig. 2 C, right panel).

**Figure 2.**
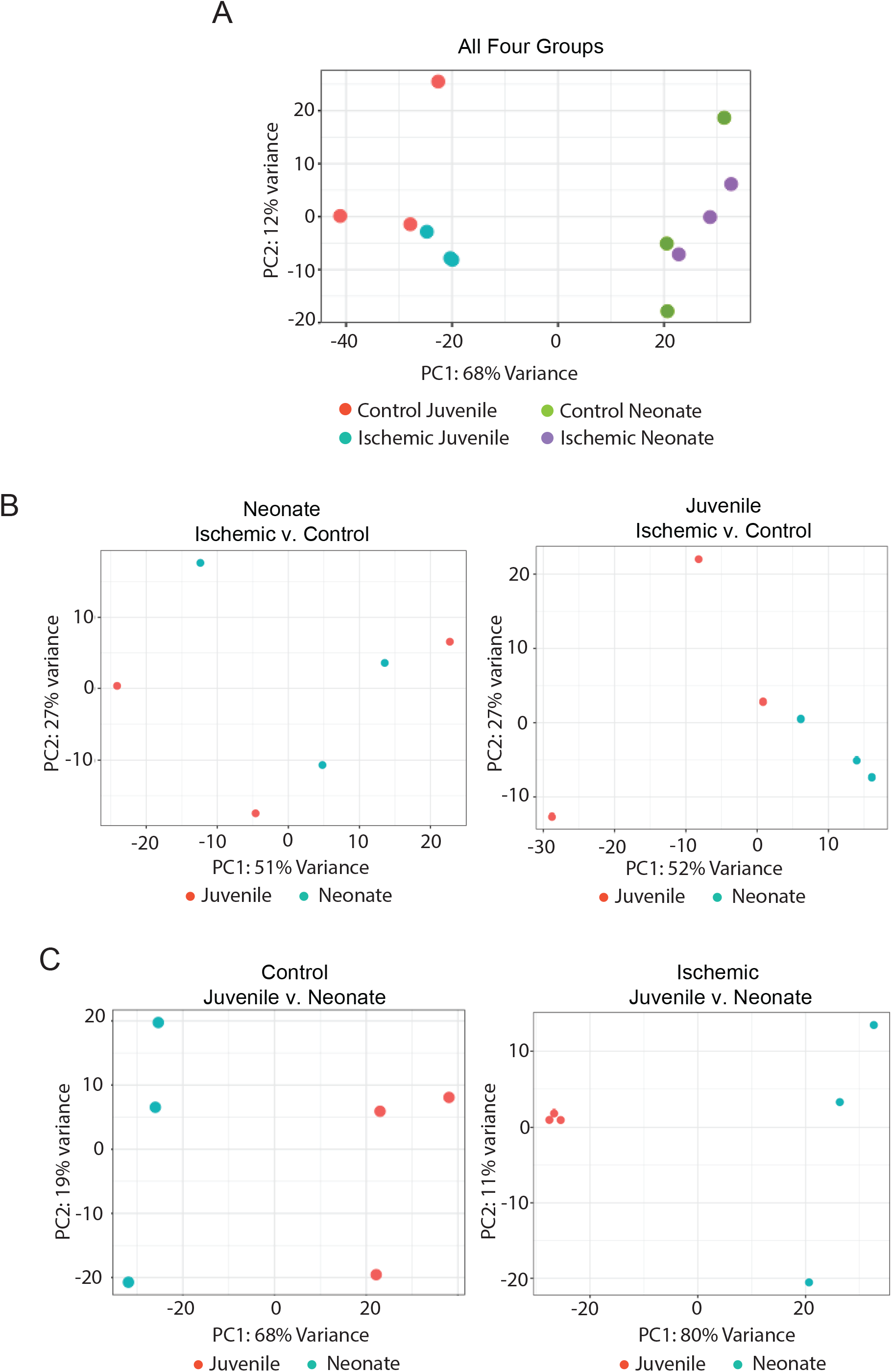
Principal components analyses (PCAs) of differential mRNA expression data identify distinct clustering in response to ischemic injury in juveniles, but not neonates. (A) PCA plots of differentially expressed genes compared across ischemia-injured or control jejunal mucosa in neonatal and juvenile pigs. Neonates (green and purple) and juveniles (red and blue) cluster distinctly across principal component 1 (PC1) with 68% of variance represented along that axis. Ischemic juveniles (blue) cluster more distinctly from control juveniles (red) than do ischemic neonates (purple) from control neonates (green). (B) When comparing ischemia effects within age groups, in neonates, there was no distinct clustering between control (red) and ischemic (blue) mucosa (left panel). Within juveniles, there was clustering of ischemic mucosa (blue) from control mucosa (red) across PC1 which contained 52% of variance (middle panel). (C) When comparing age effects within either control or ischemic groups, both control and ischemic juvenile mucosa (red) clustered separately from neonatal mucosa (blue) with 61% and 80% of variance across PC1, respectively.

Further analysis of differentially expressed genes using Ingenuity Pathways Analysis predicted downregulation of eleven “Diseases and Functions” signaling pathways instrumental in cell migration under ischemic conditions in neonatal mucosa as compared to juveniles (Table 1).

**Table 1.** **“Diseases and Functions” Ingenuity Pathways Analysis of differentially expressed gene dataset in ischemic neonatal mucosa relative to ischemic juvenile mucosa.**

| Function | P-value | z-score | Predicted State* |
| --- | --- | --- | --- |
| Cell movement | $2.26 \times 10^{-45}$ | -2.619 | ▼ |
| Migration of cells | $5.10 \times 10^{-44}$ | -2.558 | ▼ |
| Activation of cells | $1.38 \times 10^{-27}$ | -2.915 | ▼ |
| Organization of cytoskeleton | $3.72 \times 10^{-26}$ | -3.067 | ▼ |
| Cellular homeostasis | $2.44 \times 10^{-24}$ | -3.014 | ▼ |
| Microtubule dynamics | $1.84 \times 10^{-22}$ | -2.543 | ▼ |
| Organization of cytoplasm | $8.67 \times 10^{-22}$ | -3.067 | ▼ |
| Cell-cell contact | $6.26 \times 10^{-14}$ | -3.287 | ▼ |
| Formation of cell protrusions | $8.23 \times 10^{-14}$ | -2.825 | ▼ |
| Formation of filaments | $2.27 \times 10^{-13}$ | -2.498 | ▼ |
| Formation of cytoskeleton | $3.89 \times 10^{-13}$ | -3.013 | ▼ |

Finally, we used Gene Set Enrichment Analysis (GSEA) to evaluate whether our experimental groups showed significant positive or negative relative enrichment for several Molecular Signatures Database “hallmark” gene sets associated with epithelial cell functions that are relevant to epithelial wound healing. Within neonatal mucosa, the hallmark gene sets “epithelial to mesenchymal transition,” “apical junction” and “integrin-cell surface interactions” were negatively correlated with the ischemic condition (Fig. 3 B, left panels). Within juvenile mucosa, the opposite correlation was found with two of these gene sets, in that “epithelial to mesenchymal transition” and “integrin-cell surface interactions” were positively correlated with the ischemic condition (Fig. 3 B, middle panels). When comparing the ischemic condition across age groups, “epithelial to mesenchymal transition,” “apical junction” and “integrin-cell surface interactions” were negatively correlated with the neonates, enriched in juveniles, consistent with the defective epithelial barrier and defective epithelial restitution observed in this age group (Figure 3 B, right panels) (19). These results agree with the cellular phenotypes observed in neonates and juveniles, further demonstrating the lack of a coordinated transcriptional response to epithelial barrier injury in neonates as compared to juveniles.

**Figure 3.**
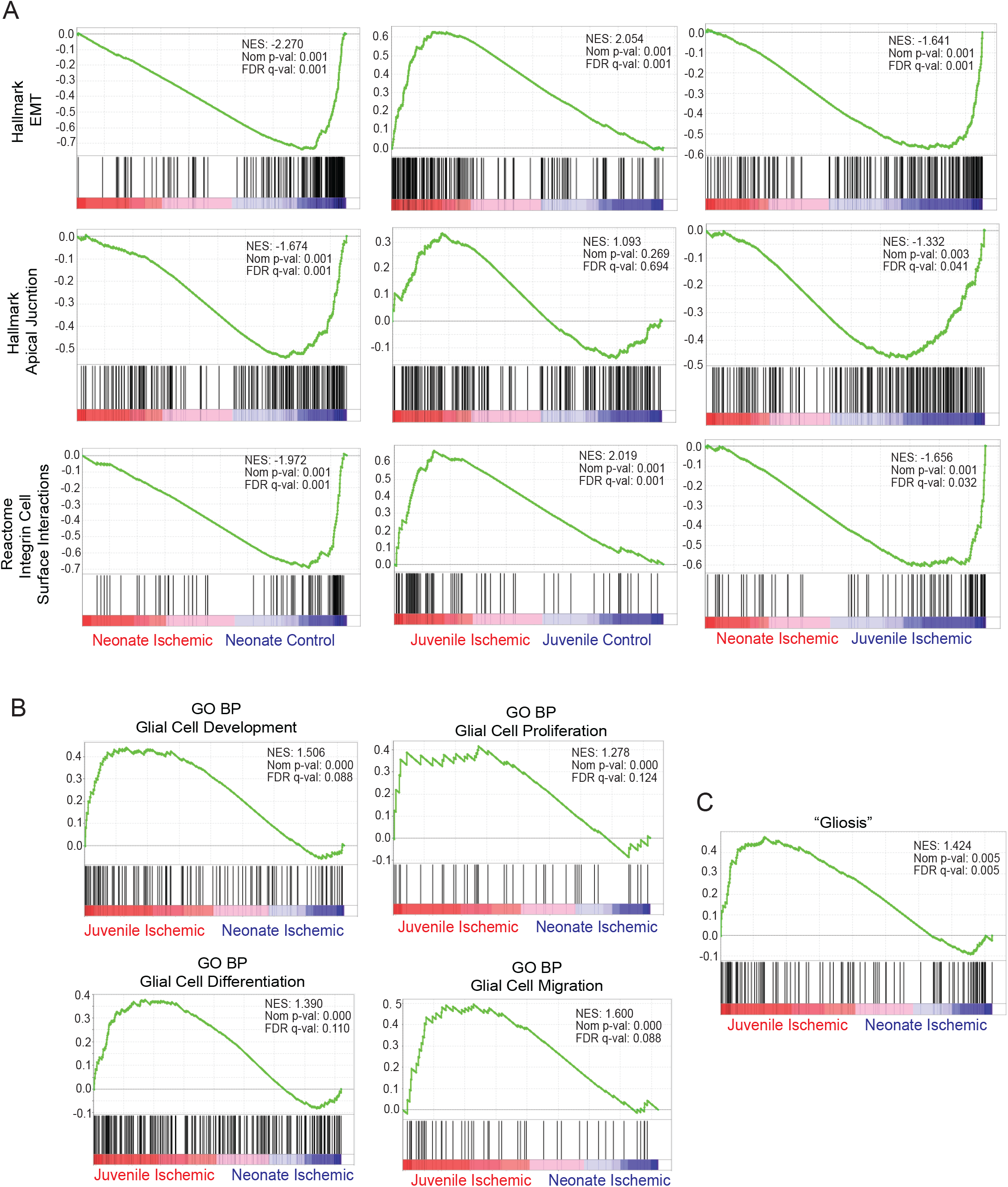
Transcriptomic differential expression analyses identify enrichment for select pathways implicated in epithelial adhesion and migration, and enteric glial cell reactivity termed “gliosis” in juvenile pigs. Gene set enrichment analyses (GSEA) of RNA-sequencing data from control and ischemia-injured neonate and juvenile mucosa. Green line shows transcripts from the specified pathway with increased (positive values) or decreased (negative values) expression by rank. Black bar coding represents location of individual genes on the green curves. Normalized enrichment score (NES), nominal *P* values and false discovery rates (FDR) *Q* values are shown on each graph. FDR *Q*-val ≥ 0.05 was considered significant. (A) Comparisons of effect of injury on transcripts within age groups (left and middle panels) and effect of age on injury transcripts (right panels) are shown. Activation of gene sets associated with epithelial migration (Hallmark “Epithelial to Mesenchymal Transition” [EMT], Hallmark “Apical Junction” and Reactome “Integrin Cell Surface Interactions” are markedly impaired in neonates (left and right columns). Within juveniles, Hallmark “Epithelial to Mesenchymal Transition” [EMT] and Reactome “Integrin Cell Surface Interactions” are enriched in ischemic mucosa (middle column). (B) Within ischemic mucosal samples, Gene Ontology Biological Processes (GO BP) “Glial Cell Development”, “Glial Cell Proliferation”, “Glial Cell Differentiation”, and “Glial Cell Migration” trend toward enrichment in juveniles as compared to neonates. (C) “Gliosis” as defined by Schneider *et al* (32) was enriched in ischemic juveniles as compared to ischemic neonates.

### Gene signatures which drive development of the enteric glial cell network and “gliosis” response to injury are enriched in ischemia-injured juvenile mucosa

Due to the predominant role of EGC in promoting epithelial restitution in adults and in light of evidence that mucosal EGC network establishes postnatally in rodents, we used GSEA to compare relative enrichment for gene datasets related to glial cell biology in transcriptomic profiles of ischemic mucosa from juvenile versus neonates.(20, 22, 23, 28) Using GSEA, we found that Gene Ontology Network Biological Processes gene sets for “glial cell development,” “glial cell proliferation,” “glial cell differentiation” and “glial cell migration” demonstrated a strong trend toward enrichment in the juvenile mucosa as compared to the neonatal mucosa (Fig. 3 B). In many disease states, including ischemia, a EGC may transition to an activated immune phenotype termed “gliosis”. This is characterized by an increased expression of a set of genes which include a subset of inflammatory response genes, and this gene set was defined in a new GO term published by Schneider *et al* in 2021.(32) We interrogated this gene set in ischemic neonatal and juvenile mucosal tissue using GSEA and, importantly, we found that “gliosis” is enriched in juvenile tissues undergoing ischemic injury as compared to neonates (Fig. 3 C).

Altogether, these transcriptomic analyses indicate that gene sets which drive development of the enteric glial cell network and “gliosis” response to injury are enriched in ischemia-injured juvenile mucosa, and the relative lack of these gene signatures in neonatal mucosa suggests that reduced EGC network development and activity might play an important role in age-dependent defects in epithelial restitution.

### The three-dimensional density of EGC within the sub-epithelial space shifts discernably during early postnatal development

In light of our transcriptomic results implicating the EGC network, we next sought to define age-dependent differences in the EGC network density in the porcine jejunal mucosa. To completely visualize and quantify changes in the three-dimensional network of EGC in the postnatal pig jejunum, we utilized a whole-tissue imaging approach called immunolabeled three-dimensional imaging of solvent cleared organs (iDISCO) in which optically cleared full-thickness sections of intestine are stained with fluorescent antibodies and imaged in three dimensions with a light sheet microscope(36). Full thickness jejunum samples stained against glial fibrillary acidic protein (GFAP), a component of the cytoskeleton in certain EGC subtypes, (37) demonstrated a dense and complex network of GFAP^+^ EGC throughout the submucosal plexus (SMP, Fig. 4 A) and longitudinal muscle myenteric plexus (LMMP, Fig. 4 A) in both neonates and juveniles. In the lamina propria, however, there was a greater density of GFAP^+^ EGC in the immediate subepithelial space of jejunal villi in neonates as compared to juveniles (Fig. 4 B, left panels). GFAP^+^ EGC were identified by a customized surfaces algorithm in Imaris^®^ software (Fig. 4 B, right panels) and quantification demonstrated that maximum GFAP signal intensity and GFAP^+^ EGC density, measured as the percent volume of the villi occupied by GFAP^+^ EGC, were increased in juvenile lamina propria compared to neonates (Fig. 4 D, E).

**Figure 4.**
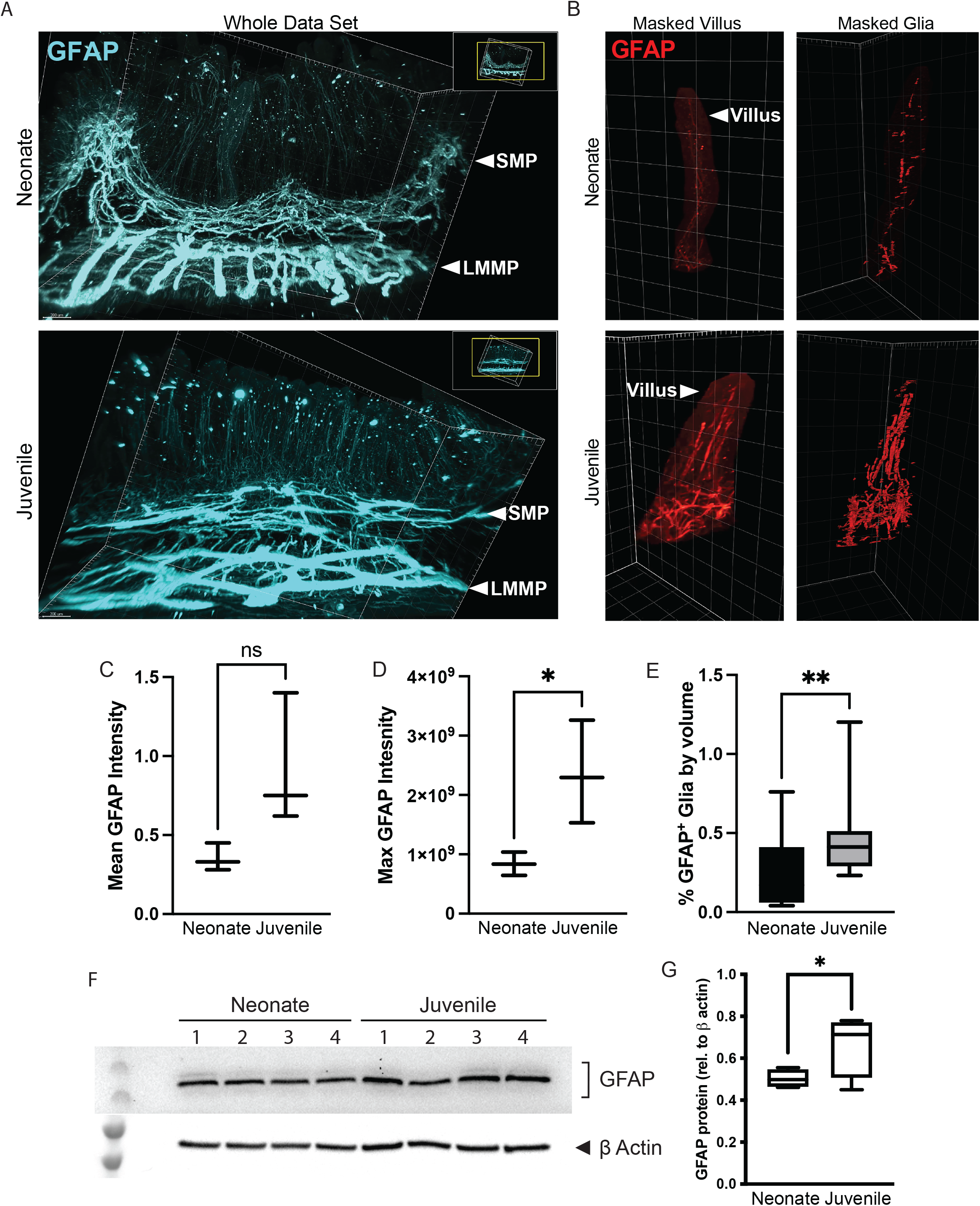
GFAP^+^ EGC density is increased in the subepithelial space of juvenile mucosa as compared to neonates. (A) Light sheet imaging of GFAP^+^ EGC networks in iDISCO-prepared whole jejunum revealed lower glial density in the lamina propria of neonatal jejunum as compared to juveniles. 2D representations of 3D EGC networks labeled against glial fibrillary acidic protein (GFAP) in neonatal (top panels) and juvenile (bottom panels) jejunum. Panels in blue represent the entire data set while panels in red represent a single villus masked (middle panels) for EGC volume quantification by custom-designed algorithm using Imaris^®^ software (right panels). (C,D) trending increased mean and increased maximum GFAP signal intensity was measured within the lamina propria of juveniles as compared to neonates. (E) Percent of villus volume occupied by GFAP^+^ glia is higher in juvenile versus neonates. (n=3-4. *P≤0.05 by one-tailed Mann-Whitney t-test) (F) GFAP was detected in a prominent 57kD band in microdissected mucosal tissue. (G) Increased GFAP protein expression was quantified in the subepithelial mucosa of juveniles as compared to neonates (n=4. *P<0.05, unpaired student’s t-test).

To further quantify changes in EGC density within the discrete layers of the jejunum during the postnatal period, we evaluated the levels of expression of GFAP in jejunal mucosa isolated from neonatal and juvenile pigs by microdissection. GFAP signal within the mucosa was restricted to primarily one band at 57kD (Fig. 4 F). There was increased GFAP in juveniles as compared to neonates (Fig. 4 G), in agreement with transcriptomic trends (Fig. 3 B) and 3D immunofluorescence quantification (Fig. 4 D, E). Altogether, these results suggest that the three-dimensional distribution of GFAP^+^ EGC into the immediate subepithelial space is enhanced in juvenile-aged jejunal mucosa as compared to neonates.

### Fluoroacetate inhibits barrier restitution after ischemic injury in juveniles

The gliotoxin fluoroacetate (FA) has been used experimentally to inhibit glial cell metabolism and activity, therefore we sought to utilize FA to inhibit glial cell activity to directly implicate of the EGC network on barrier restitution.(38-40) To examine whether FA-mediated inhibition of EGC would induce a restitution defect in the repair-competent juvenile pig model, barrier function was examined by both electrophysiology and histomorphometry during *ex vivo* recovery from 30-minutes of intestinal ischemia with and without the application of increasing doses of FA. Treatment with any dose of FA during *ex vivo* recovery inhibited recovery of TEER in juvenile jejunal mucosa injured by 30-minutes of surgical ischemia (Fig. 5 A). Histomorphometry revealed significant loss of epithelium at a high dilution of 5000μM FA even in the absence of ischemic injury (Fig. 5 B, green bars). At a low dilution of 50μM FA, epithelial coverage was similar to control tissues across all combinations of ischemia and recovery (Fig. 5 B, yellow and black bars, respectively). At a mid-range dilution of 500μM FA, there was no significant loss of epithelium in the absence of ischemic injury, however recovery of epithelial coverage during *ex vivo* recovery after ischemia injury was inhibited, recreating the restitution defect seen in the neonatal model (Fig. 5 B, blue bars). Histological appearance of non-ischemic mucosal epithelium appeared very similar after 120 minutes *ex vivo* incubation in Ussing chambers with and without treatment with 500μM FA. Complete epithelialization was observed at the mucosal surface of villi treated with 500μM FA (Fig. 5 C, open arrowheads) providing evidence of a lack of a non-specific toxic effect of FA at this dosage on epithelial morphology in uninjured jejunum. Following 30-minutes intestinal ischemic injury, tissues incubated *ex vivo* in control conditions demonstrated effective restitution characterized by the flatted morphology of the newly restituted epithelial cells at the villus tips as previously reported, contrasting with the persistent epithelial defects observed in the tissues incubated *ex vivo* in 500μM FA, which demonstrated a persistent epithelial defect at the tip of the villi, very similar to previously reported restitution defect in neonates (Fig. 5 C, closed arrowheads)(19). Along with other phenotypical parameters, elevated levels of GFAP in EGC are indicative of “gliosis” in some disease states, including ischemia. (32, 41-44). To validate that 500μM FA treatment suppresses activation of GFAP expression in EGC in response to ischemic injury in our model, we measured FA-mediated inhibition of ischemia-induced increases in GFAP signal intensity in the submucosal EGC (Fig. S1). Altogether, these results suggest that EGC are instrumental in orchestrating mucosal restitution in ischemia-injured jejunum of juvenile pigs.

**Figure 5.**
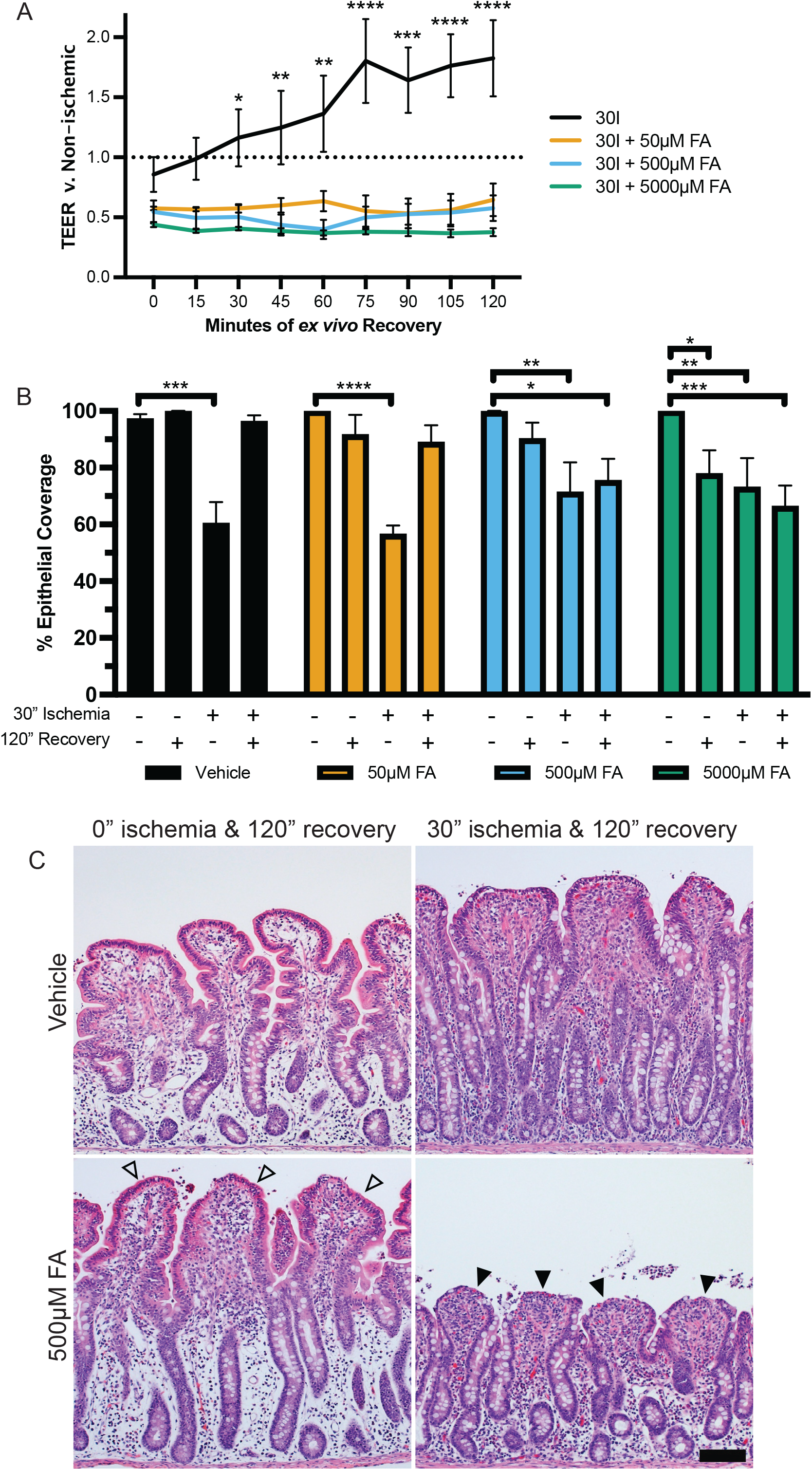
Treatment with 500μM EGC inhibitor fluoroacetate (FA) inhibits restitution in ischemia-injured juvenile jejunum. (A) No recovery of TEER was seen in the presence of FA, regardless of concentration. (n=3-4. P<0.0001 for overall effect of treatment on TEER over time by two-way ANOVA. *P<0.05, **P<0.01, ***P<0.001, ****P<0.0001 by Dunnett’s multiple comparisons test.) (B) 50μM FA did not significantly alter epithelialization patterns versus tissues incubated in standard Ringers solution. 5mM FA inflicted significant epithelial loss in the absence of ischemic injury. Without causing significant epithelial loss in the absence of ischemic injury, 500μM FA inhibited epithelial restitution following ischemia. (n=3-4. P<0.0001 for overall effect of ischemia condition on TEER over time and a significant interaction between ischemia condition and treatment [P=0.0223] by two-way ANOVA. *P<0.05, **P<0.01, ***P<0.001, ****P<0.0001 by Dunnett’s multiple comparisons test. Scale bar = 50μm.) C. Representative micrographs showing effect of 500μM Fluoroacetate on epithelial restitution. Note no significant epithelial injury in non-ischemic mucosa (open arrowhead) but inhibited restitution in ischemic mucosa (solid arrowhead).

### Co-culture with neonatal EGC enhances *in vitro* restitution in intestinal epithelial cells

To examine glial-epithelial interactions more precisely during barrier repair, a co-culture model with primary pig EGC was implemented. Primary pig EGC were isolated from the submucosal (SMP) and longitudinal muscle myenteric plexuses (LMMP) of uninjured pig jejunum. Purity of culture was confirmed by the expression of the EGC markers GFAP and S100β with very few negative cells present (Fig. 6 A, open arrowheads). Representative images of primary EGC culture showed cells with fibrillar arrangement of GFAP positive cytoskeletal elements, notably in cellular projections (arrows), and a more diffuse pattern for S100β signal. For restitution assays in a transwell system, neonatal pig jejunum EGC were maintained on the bottom of a well and placed in the presence of IPEC-J2 confluent monolayers seeded on permeable membranes in the apical chamber suspended above the EGC for 24 hours to allow glial-epithelial paracrine bi-directional exchanges of soluble factors prior to manually inflicting a scratch wound (Fig. 6 B). Six hours after wounding, there was a significant increase in restitution rates in IPEC-J2 monolayers in co-culture with pig EGC from either the SMP or LMMP as compared to IPEC-J2 restitution in monoculture. Measured wound closure at 6-hours after wounding was 77.4% and 73.0% for SMP and LMMP, respectively, versus 35.6% in monoculture. These *in vitro* results provide important proof-of-concept that important pro-restitution communications may take place between the epithelium and EGC when these two cell populations are in great enough density and proximity to one another for paracrine signaling interactions to occur (Fig. 6 C).

**Figure 6.**
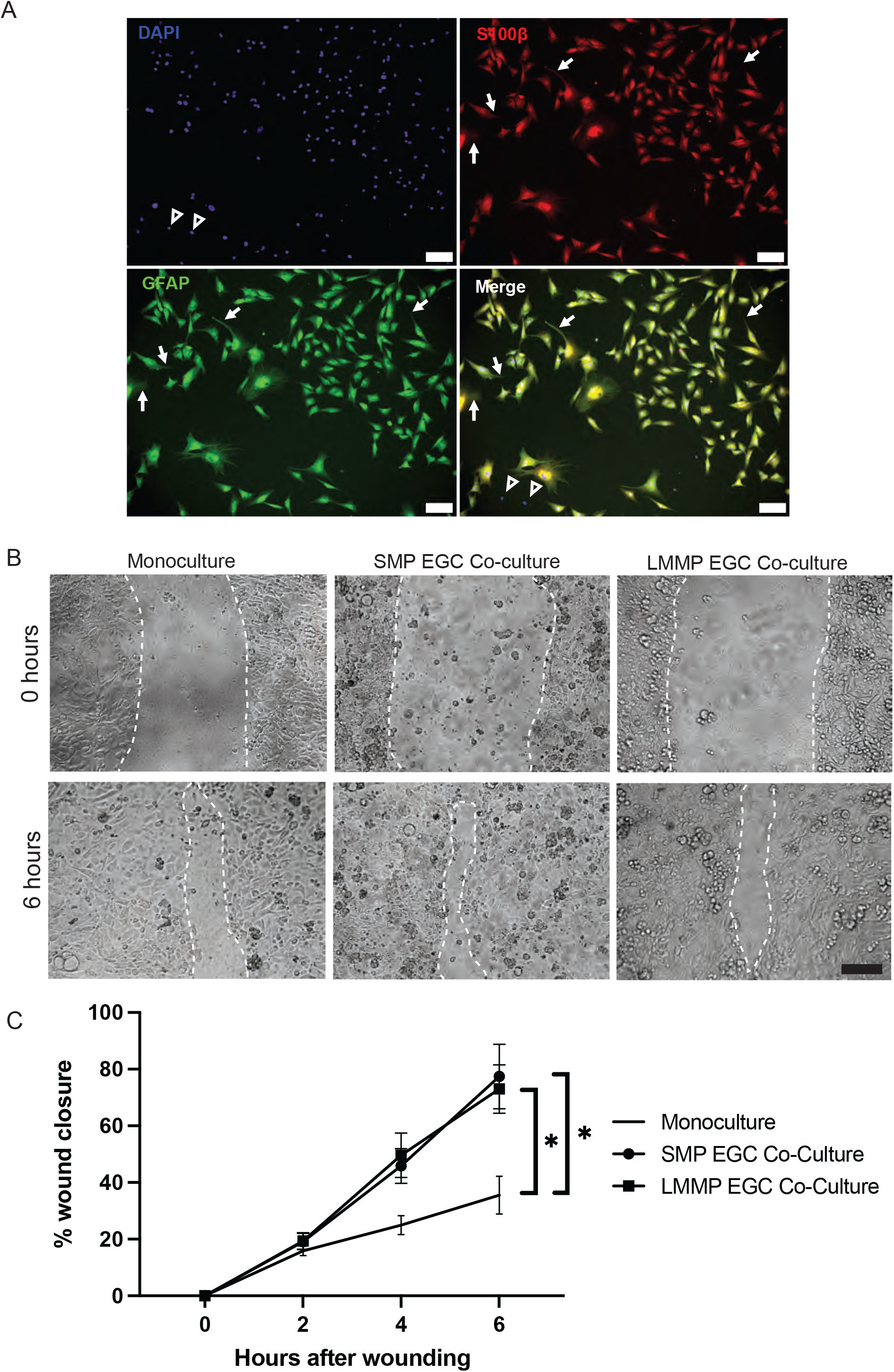
IPEC-J2 monolayers restitute scratch wounds more efficiently when in co-culture with pig EGC. (A) Pig EGC can be isolated and maintained *in vitro*. Primary EGC from pig jejunal SMP in culture immunolabeled for EGC markers S100β (red) and GFAP (green). Note the small number of negative cells (open arrowheads) and the fibrillary staining pattern of GFAP versus the cytoplasmic pattern of S100β (arrows) in the co-labeled EGC. Scale bars = 100μm. (B) Representative photomicrographs of IPEC-J2 monolayers at 0- and 6-hours after scratch wounding when in monoculture (left) or in co-culture with SMP EGC (middle) or LMMP EGC (right). Wound margins are outlined with a broken white line. Scale bars are 100μm. (C) Quantification of wound closure rates based on wound surface areas at 0-, 2-, 4- and 6-hours after wounding revealed a significant effect of time (P<0.0001) and co-culture (P=0.019) and an interaction between time and co-culture (P=0.0002) on wound healing was observed by two-way repeated measures ANOVA (n=6). Tukey’s multiple comparisons test revealed a significant difference between SMP co-culture versus monoculture (P=0.026) and between LMMP co-culture and monoculture (P=0.021) at 6-hours post wounding (*P≤0.05).

## DISCUSSION

In this study, we initially further characterized epithelial restitution after intestinal ischemic injury into later stages of acute mucosal recovery when juvenile tissues have fully recovered barrier function, but neonates have not, to improve our understanding of the epithelial repair defects observed in neonates. Scanning EM imaging confirmed juvenile mucosa exhibits the characteristic epithelial phenotype of restitution, defined by the recruitment membrane from the microvilli into the extension of both lamellipodia and filopodia onto the exposed extracellular matrix of the wound beds. This is highly indicative of successful epithelial migration and wound closure, and indeed, at later stages of acute mucosal recovery, all villi were completely covered with overlapping epithelial cells of this flattened and irregular appearance with some persisting lamellipodial extensions visible. In contrast, neonatal epithelial cells remained round and covered in microvilli throughout later stages of recovery, without any phenotypic indications of directional migration. This phenotype closely parallels what has been described in other intestinal injury models, such as a punch biopsy murine intestinal wound healing model in which the authors define “wound associated epithelial (WAE) cells” characterized by migratory nature and a shorter height than non-WAE cells, or *in vitro* scratch-wounding epithelial injury models which describe intestinal epithelial cell spreading and migrating in a similar manner.(45-47) These findings substantiate an age-dependent restitution defect defined by the striking contrast in the morphology of newly restituted epithelial cells in juvenile jejunum versus the complete absence of a restitution phenotype observed in wound-adjacent epithelial cells of neonates at later phases of acute barrier repair.

To identify key age-dependent differences in the molecular mechanisms occurring in injured jejunum, we assessed transcriptomic changes occurring in the mucosa of neonates *versus* juveniles in response to ischemic injury by bulk RNA sequencing. In agreement with the striking observed phenotypic differences, PCA plots of transcriptional expression data from uninjured and injured neonatal and juvenile mucosa suggested a lack of a coordinated transcriptional response in the neonatal mucosa in response to injury while juveniles demonstrate tight clustering in response to injury across multiple methods of PCA comparison. These age-dependent differences in coordinated transcriptional responses parallel the age-dependent response in the wound-associated epithelial cells observed by scanning EM. To identify global signaling mechanisms associated with these transcriptomic changes, we used two complementary analyses, the first one identifying pathways and mechanisms representative of the significantly differentially expressed genes (IPA) and the second one testing for the relative enrichment of pre-defined gene sets in global transcriptomic profiles (GSEA). When comparing injured neonatal mucosa to injured juvenile mucosa to interrogate solely age-dependent differences in transcriptional response to injury, several interesting pathways emerged. Two gene sets important in epithelial adhesion, Hallmark “Apical Junction” and Reactome “Integrin Cell Surface Interactions,” were found to be enriched in ischemia-injured juveniles versus ischemia-injured neonates. Apical junctional complexes contain the zonula adherens and the tight junctions, the latter of which are the molecular structures most integral to sealing the interepithelial spaces and forming the intestinal barrier and are instrumental in intestinal barrier repair.(48) Integrins are essential to epithelial cell adhesion to the basement membrane, and their precise regulation is critical to cellular migration during coordinated wound healing events in the intestine, as has been shown with integrin β1 in restituting and hypoxic intestinal epithelium.(45, 49) Integrins have been shown to also regulate the activation of various intracellular molecular mechanisms controlling cell functions key to mucosal repair, such as the role of α3β1 integrin in inhibiting Smad7, effectively potentiating TGF-β1-mediated keratinocyte migration during cutaneous wound healing.(50) Thus, alterations in integrin signaling at the RNA level suggest potential important implications in epithelial wound healing. Furthermore, pathways identified by IPA listed in Table 1 further substantiate reduced cellular migration-associated signaling in the ischemic neonatal mucosa as compared to ischemic juvenile mucosa, including regulation of cytoskeleton, microtubules, filaments, and cellular protrusions, and more broadly “cell movement” and “migration of cells”. Altogether EM and RNA-seq data strongly support previous findings demonstrating major defects in epithelial restitution following intestinal ischemia in neonatal versus juvenile pigs.(19)

Interestingly, the Hallmark Gene Set “Epithelial to Mesenchymal Transition” (EMT) was also found to be enriched in juveniles *versus* neonates after injury. This would indicate that there is an EMT-like pathway associated with acute re-epithelialization in the intestine, which is absent in the neonates. EMT is a well-described predominantly transcriptionally based trans-differentiation process that consists of coordinated phenotypic changes in epithelial cells leading notably to reduction of cell-cell adhesion, loss of cell polarity and adoption of a migratory phenotype, just as is observed in intestinal restitution following ischemic injury in juveniles. This transition has been defined in embryogenesis (type 1 EMT), inflammation and organ fibrosis (type 2 EMT) and epithelial origin cancers (type 3 EMT).(51) Most notably, EMT is a hallmark of metastatic neoplastic disease of epithelial origin as this cellular phenotype is highly mobile.(52) An intestinal epithelial cell’s transition to a non-neoplastic, non-fibrotic migratory state during acute barrier repair may involve very similar signaling pathways as EMT.

As we have previously reported, the neonatal defect in restitution is rescued by direct application of ischemia-injured jejunal homogenate from juvenile-aged pigs, but the component of this homogenate which drives restitution has not yet been identified(19). Given the emerging and growing appreciation for the critical role of EGC in regulating and relaying signaling events in the intestinal mucosa in both homeostatic and pathologic conditions, we have considered their influence in our model.(53-55) Of particular relevance to our studies, many have demonstrated direct interactions between EGC and the intestinal epithelium to direct the maintenance and repair of the intestinal barrier.(23, 26-28, 56-62) In a 2009 study, co-culture with transformed rat EGC (CRL2690 cell line) markedly altered transcriptomic regulation of pathways relevant to intestinal barrier function in Caco-2 colon cancer epithelial cells. More specifically, microarray analysis demonstrated that EGC increase transcriptional pathways related to cell adhesion, differentiation, and motility in intestinal epithelial cells –thereby favoring enhanced epithelial barrier function and healing.(25) In fact, additional functional studies have provided strong evidence using *in vitro* systems and *in vivo* rodent models that EGC promote barrier function and epithelial restitution after injury in adulthood. (23, 26) Therefore, it was tempting to speculate that the molecule(s) present in jejunal homogenate from ischemia-injured juvenile pigs and responsible for rescuing restitution in neonates originated from EGC. Supporting this hypothesis, prevailing evidence in rodents suggests that the EGC network continues maturation after birth leading us to surmise that these networks may be underdeveloped in the neonatal condition in the pig model (22, 63, 64). Interestingly lineage tracing experiments have elegantly demonstrated that mucosal EGC network establishes from EGC migrating along a serosa-mucosa axis, *i*.*e*., originating from the immediate underneath layer, the submucosal plexus, and this seems to be also true for the pig model.(22, 65). Indeed, GSEA demonstrated a strong trend toward enrichment of molecular pathways involved in glial cell development and migration in the juvenile *versus* neonatal ischemic mucosa. This would suggest that pathways coordinating EGC migration and network development into the mucosa are more active in juvenile mucosa as compared to neonates, which is consistent with a putative role played by the EGC network in the age-dependent defects in epithelial restitution observed during the postnatal period. In fact, this was supported further by three-dimensional volume imaging and western blot analysis quantifying a definite increase in GFAP^+^ EGC density within the subepithelial region of juvenile pig jejunum as compared to neonates. Altogether, these findings are indicative of increased GFAP^+^ EGC density in the subepithelial space, which may reflect increased numbers of GFAP^+^ cells, or increased expression of GFAP in individual cells. These subepithelial EGC are the population most relevant to paracrine regulation of epithelial barrier as paracrine signaling requires an optimal distance of just a few microns between communicating cells.(66) Of note, EGC of different phenotypes have been shown to differentially express common EGC markers; at least four described subtypes have been proposed without any single definitive pan-EGC marker yet described.(37) The striking heterogeneity of EGC as well as the specific locations of these distinct subtypes is increasingly suspected to result in EGC subtype-specific impact on epithelial cell functions.(67) Efforts to fully map postnatal changes in EGC network density and cellular composition within the sub-epithelial space is an important area for ongoing study.

Further substantiating our notion that EGC activity may be responsible for age-dependent failure of restitution, induction of “gliosis” as defined by Schneider *et al* in 2021 was found to be diminished in neonatal mucosa during ischemic injury as compared to juveniles. In the central nervous system, gliosis is a nonspecific, universal response of glia to injury, and results from many acute conditions including ischemia and stroke.(68) In the gut, gliosis involves enhanced inflammatory signaling in EGC, and it has been reported increasingly in the literature in response to many intestinal disease and injury states like chemotherapy, inflammation, infection and motility disorders.(32, 57, 69-72) This reactive state is associated with the release of many cytokines and glial factors which would have direct interactions with the mucosal barrier, and thus could be an important component to mucosal response to acute injury as seen in this pig model of acute intestinal injury. In keeping with this, when treating injured EGC with glial cell inhibitor fluoroacetate (FA), ischemia-induced upregulation of GFAP, one described marker of the EGC gliosis phenotype, is inhibited.(32, 38) Taken together with the changes observed in EGC network development in the immediate subepithelial space, these findings demonstrate clear age-dependent differences in the EGC populations of the neonatal and juvenile jejunum which can carry important implications in this model.

Although strong evidence exists using *in vitro* co-culture systems and *in vivo* murine models, to the best of our knowledge this study is the first to report that EGC promote epithelial restitution after ischemia in a large animal model. Indeed, using FA to inhibit EGC activity in juvenile mucosa, we confirmed that EGC inhibition abrogates barrier function recovery and epithelial restitution in juvenile mucosa recovering from 30-minutes ischemic injury, mirroring the neonatal restitution defect described by our lab in 2018.(19) In a complementary approach and to demonstrate proof-of-concept more fully, an *in vitro* scratch wound model of intestinal barrier injury and repair using co-culture of a neonatal pig intestinal epithelial cell monolayer with jejunal EGC from either the SMP or LMMP of neonatal pigs demonstrated two-fold enhanced restitution efficiency. Undeniably, neonatal EGC can enhance epithelial barrier restitution *in vitro* by secreted factors which readily diffuse through cell culture media. Feasibly, this indicates an efficient pro-repair paracrine signaling mechanism is present in neonatal EGC which are in the deeper layers of the jejunum, but that these cells are too sparse in the immediate sub-epithelial space to efficiently stimulate epithelial restitution during this early postnatal period. Seemingly, this signaling derived from neonatal EGC can be effective in driving intestinal epithelial repair once the EGC density near wounded epithelium is sufficient as it was in this reductive culture system. Existing reported findings predicted and support these results; EGC are known to regulate intestinal epithelial barrier in transformed cell lines *in vitro*, and more specifically, a paracrine effect of ECG in driving intestinal barrier restitution *in vitro* has been previously demonstrated in wounded Caco2 cells, and was found to be mediated by proEGF driving FAK-mediated epithelial restitution, which may be an important pathway to investigate in ongoing studies.(23, 25) These data presented here are the first to demonstrate this effect with porcine intestinal epithelial cells, showing promise for high translation potential in complex large animal models. Ongoing work to understand the EGC-mediated paracrine signaling *ex vivo* and *in vivo* in highly comparative models is essential to driving translational efforts to improve clinical recovery of neonatal patients with intestinal ischemic injury.

Altogether, our results further refine our model of postnatal age-dependent defects in epithelial repair characterized by a failure in neonates to activate both transcriptional and ultrastructural epithelial restitution phenotypes following ischemia. Importantly, they provide key evidence that a dense network of GFAP^+^ EGC near the damaged epithelium play a critical role in promoting epithelial barrier repair in juveniles, and that a sub-optimal density of a mature subepithelial EGC network in neonates is likely contributing to the observed age-dependent defect in epithelial barrier restitution. Future studies should seek to understand the specific ways in which the mucosal microenvironment, and specifically EGC, are modulated during the postnatal period to promote barrier repair. Novel therapeutic strategies targeting specific epithelial repair signaling mechanisms identified in this highly comparative pig model have great potential to improve outcomes in neonates suffering from intestinal failure associated with epithelial barrier disruption.

## ACKNOWLEDGEMENTS

The authors would like to thank Lindsey Shapiro and Kayla Harrower for their assistance with tissue collection and sample preparations, as well as quantitative analyses.

## GRANTS

NIH K01 OD 028207 (ALZ); NIH-NICHD R01 HD095876 (ATB, JO, LVL); USDA-NIFA 2019-67017-29372 (ATB, JO, LVL); P30 DK034987 (ATB)

## DISCLOSURES

The authors declare no perceived or potential conflict of interest, financial or otherwise.

## FIGURE CAPTIONS

**Supplementary Figure 1.**
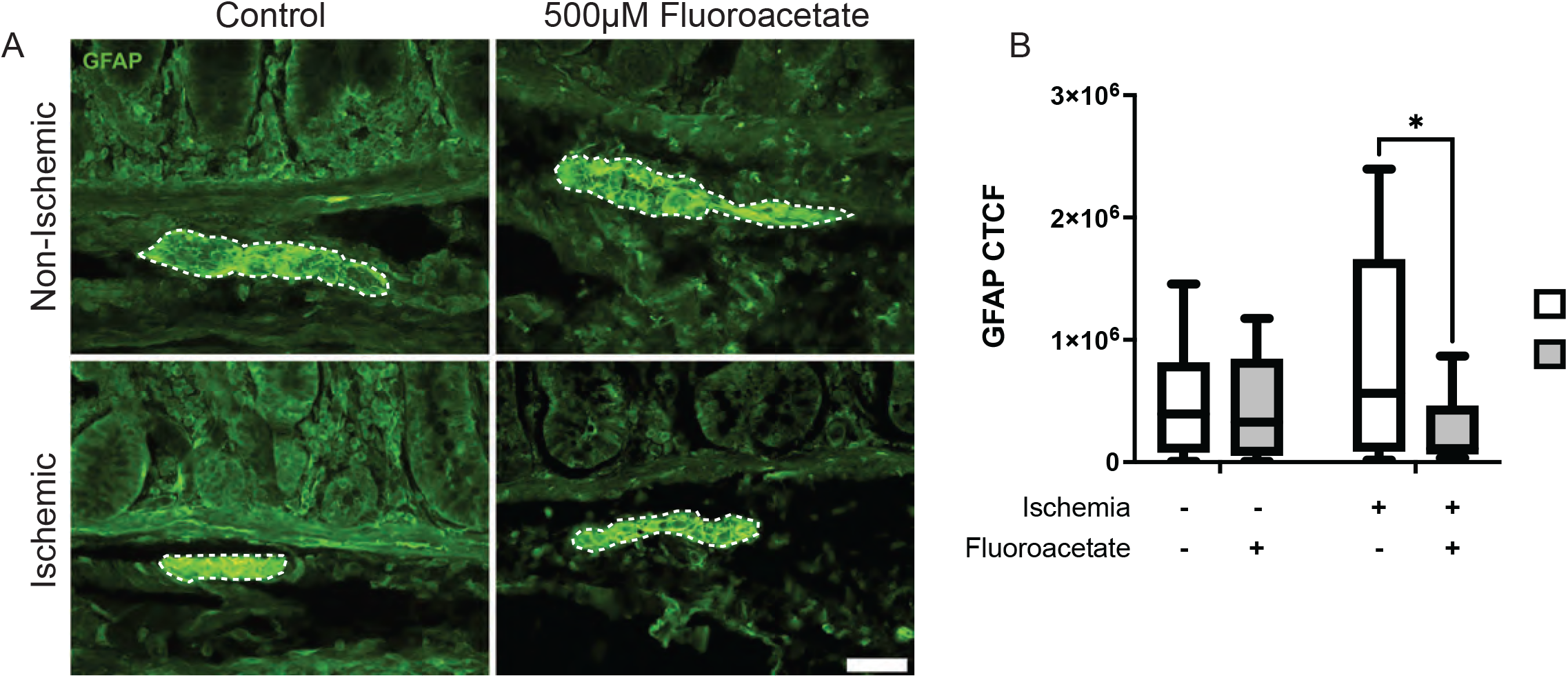
500μM fluoroacetate suppresses increased GFAP intensity in response to ischemic injury. Boxplot (min to max) distribution depicting GFAP corrected total cell fluorescence intensity expressed by submucosal EGC from juvenile pig jejunum exposed to either 0 minutes of ischemia and 0 minutes of recovery (0I0R) or 30 minutes of ischemia and 0 minutes of recovery (30I0R) and treated with either 0 or 500μM fluoroacetate (FA). EGC from ischemia-injured tissue treated with 500μM showed significantly lower GFAP signal intensity than in EGC from untreated ischemia-injured tissue. (P=0.010 by Wilcoxon matched-pairs signed-rank test with Holm-Šidák method, n=3 pigs per group, 5 ganglia measured per pig).

## Notes

https://www.ncbi.nlm.nih.gov/geo/query/acc.cgi?acc=GSE212533

### Competing Interest Statement

The authors have declared no competing interest.

